# Interactive Visual Exploration and Refinement of Cluster Assignments

**DOI:** 10.1101/123844

**Authors:** Michael Kern, Alexander Lex, Nils Gehlenborg, Chris R. Johnson

## Abstract

**Background:** With ever-increasing amounts of data produced in biology research, scientists are in need of efficient data analysis methods. Cluster analysis, combined with visualization of the results, is one such method that can be used to make sense of large data volumes. At the same time, cluster analysis is known to be imperfect and depends on the choice of algorithms, parameters, and distance measures. Most clustering algorithms don’t properly account for ambiguity in the source data, as records are often assigned to discrete clusters, even if an assignment is unclear. While there are metrics and visualization techniques that allow analysts to compare clusterings or to judge cluster quality, there is no comprehensive method that allows analysts to evaluate, compare, and refine cluster assignments based on the source data, derived scores, and contextual data.

**Results:** In this paper, we introduce a method that explicitly visualizes the quality of cluster assignments, allows comparisons of clustering results and enables analysts to manually curate and refine cluster assignments. Our methods are applicable to matrix data clustered with partitional, hierarchical, and fuzzy clustering algorithms. Furthermore, we enable analysts to explore clustering results in context of other data, for example, to observe whether a clustering of genomic data results in a meaningful differentiation in phenotypes.

**Conclusions:** Our methods are integrated into Caleydo StratomeX, a popular, web-based, disease subtype analysis tool. We show in a usage scenario that our approach can reveal ambiguities in cluster assignments and produce improved clusterings that better differentiate genotypes and phenotypes.

## Introduction

Rapid improvement of data acquisition technologies and the fast growth of data collections in the biological sciences increase the need for advanced analysis methods and tools to extract meaningful information from the data. Cluster analysis is a method that can help make sense of large data and has played an important role in data mining for many years. Its purpose is to divide large datasets into meaningful subsets (clusters) of elements. The clusters then can be used for aggregation, ordering, or, in biology, to describe samples in terms of subtypes and to derive biomarkers. Clustering is ubiquitous in biological data analysis and applied to gene expression, copy number, and epigenetic data, as well as biological networks or text documents, to name just a few application areas.

A cluster is a group of similar items, where similarity is based on comparing data items using a measure of similarity. Cluster analysis is part of the standard toolbox for biology researchers, and there is a myriad of different algorithms designed for various purposes and with differing strengths and weaknesses. For example, clustering can be used to identify functionally related genes based on gene expression, or to categorize samples into disease subtypes. Since Eisen et al. [1] introduced cluster analysis for gene expression in 1998, it has been widely used to classify both, genes and samples in a variety of biological datasets [2, 3, 4, 5, 6].

However, while clustering can be useful, it is not always simple to use. Scientists have to deal with several challenges: the choice of an algorithm for a particular dataset, the parameters for these algorithms (e.g., the number of expected clusters), and the choice of a suitable similarity metric. All of these choices depend on the dataset and on the goals of the analysis. Also, methods generally suitable for a dataset can be sensitive to noise and outliers in the data and produce poor results for a high number of dimensions.

Several (semi)automated cluster validation, optimization, and evaluation techniques have been introduced to address the basic challenges of clustering and to determine the amount of concordance among certain outcomes (e.g., [7, 8, 9]). These methods try to examine the robustness of clustering results and guess the actual number of clusters. This task is often accompanied by visualizations of these measures by histograms or line graphs. Consensus clustering [10] addresses the task of detecting the number of clusters and attaining confidence in cluster assignments. It applies clustering algorithms to multiple perturbed subsamples of datasets and computes a consensus and correlation matrix from these results to measure concordance among them, and explores the stability of different techniques. These matrices are plotted both as histograms and two-dimensional graphs to assist scientists in the examination process.

Although cluster validation is a useful method to examine clustering algorithms it does not guarantee to reconstruct the actual or desired number of clusters from each data type. In particular, cluster validation is not able to compensate weaknesses of cluster algorithms to create an appropriate solution if the clustering algorithm is not suitable for a given dataset.

While knowledge about clustering algorithms and their strengths and weaknesses, as well as automated validation methods are helpful in picking a good initial configuration, trying out various algorithms and parametrizations is critical in the analysis process. For that reason, scientists usually conduct multiple runs of clustering algorithms with different parameters and compare the varying results while examining the concordance or discordance among them.

In this paper we introduce methods to evaluate and compare clustering results. We focus on revealing specificity or ambiguity of cluster assignments and embed our contributions in StratomeX [11, 12], a framework for stratification and disease subtype analysis that is also well suited to cluster comparison. Furthermore, we enable analysts to manually refine clusters and the underlying cluster assignments to improve ambiguous clusters. They can transfer entities to better fit clusters, merge similar clusters, and exclude groups of elements assumed to be outliers. An important aspect of this interactive process is that these operations can be informed by considering data that was not used to run the clustering: when considering cluster refinements, we can immediately show the impact on, for example, survival data.

In our tool, users are able to conduct multiple runs of clustering algorithms with full control over parametrization and examine both conspicuous patterns in heatmaps and quantify the quality and confidence of cluster assignments simultaneously. Our measures of cluster fit are independent from the underlying stratification/clustering technique and allows investigators to set thresholds to classify parts of a cluster as either reliable, uncertain, or a bad fit. We apply our methods to matrices of genomic datasets, which covers a large and important class of datasets and clustering applications.

We evaluate our tool based on a case study with gene expression data from The Cancer Genome Atlas and demonstrate how visual inspection and manual refinement can be used to identify new clusters.

## Background

In this section we briefly introduce clustering algorithms and their properties, as well as StratomeX, the framework we used and extended for this this research.

### Cluster Analysis

Clustering algorithms assign data to groups of similar elements. The two most common classes of algorithms are partitional and hierarchical clustering algorithms [13]; less frequently used are probabilistic or fuzzy clustering algorithms.

**Partitional algorithms** decompose data into nonoverlapping partitions that optimize a distance function, for example by reducing the sum of squared error metric with respect to Euclidean distance. Based on that, they either attempt to iteratively create a user-specified number of clusters, like in k-Means [14] or they utilize advanced methods to guess the number of clusters implicitly, such as Affinity Propagation [15].

In contrast to that, **hierarchical clustering** algorithms generate a tree of similar records by either merging smaller clusters into larger ones (agglomerative approach) or splitting groups into smaller clusters (divisive). In the resulting binary tree, commonly represented with a dendrogram, each leaf node represents a record, each inner node represents a cluster as the union of its children. Inner nodes commonly also store a measure of similarity among their children. By cutting the tree at a threshold, we are able to obtain discrete clusters from the similarity tree.

These approaches use a deterministic cluster assignment, i.e., elements are assigned exclusively to one cluster and are not in other clusters. In contrast, **fuzzy clustering** uses a probabilistic assignment approach and allows entities to belong to multiple clusters. The degree of membership is described by weights, with values between 0 (no membership at all) and 1 (unique membership to one cluster). These weights are commonly called probabilities capturing the likelihood of an element belonging to a certain partition. A prominent example algorithm is Fuzzy c-Means [16].

Clustering algorithms make use of a measure of similarity or dissimilarity between pairs of elements. They aim to maximize pair-wise similarity or minimize pairwise dissimilarity by using either geometrical distances or correlation measures. A popular way to define similarity is a measure of geometric distance based on, for example, squared Euclidean or Manhattan distance. These measures work well for “spherical” and “isolated” groups in the data [17] but are less well suited for other shapes and overlapping clusters. More sophisticated methods measure the cross-correlation or statistical relationship between two vectors. They compute correlation coefficients that denote the type of concordance and dependence among pairs of elements. The coefficients range from -1 (opposite or negative correlation) to 1 (perfect or positive correlation), whereas zero values denote that there is no relationship between two elements. The most commonly used coefficient in that context is the Pearson product-moment correlation coefficient that measures the linear relationship by means of the covariance of two variables. Spearman’s rank correlation coefficient is another approach to estimate concordance similar to Pearson’s but uses ranks or scores for data to compute covariances.

The choice of distance measure has an important impact on the clustering results, as it drives an algorithm’s determination of similarity between elements. At the same time, we can also use distance measures to identify the fit of an element to a cluster, by, for example, measuring the distance of an element to the cluster centroid. In doing so, we do not necessarily need to use the same measure that was used for the clustering in the first place. In our technique, we visualize this information for all elements in a cluster, to communicate the quality of fit to a cluster.

### StratomeX

StratomeX is a visual analysis tool for the analysis of correlations of stratifications [11, 12]. This is especially important when investigating disease subtypes that are believed to have a genomic underpinning. Originally developed as a desktop software tool, it has since been ported to a web-based client-server system [18]. Figure 1 shows an example of the latest version of StratomeX. By integrating our methods into StratomeX, we can also consider the relationships of clusters to other datasets, including clinical data, mutations, and copy number alteration of individual genes.

**Figure 1.**
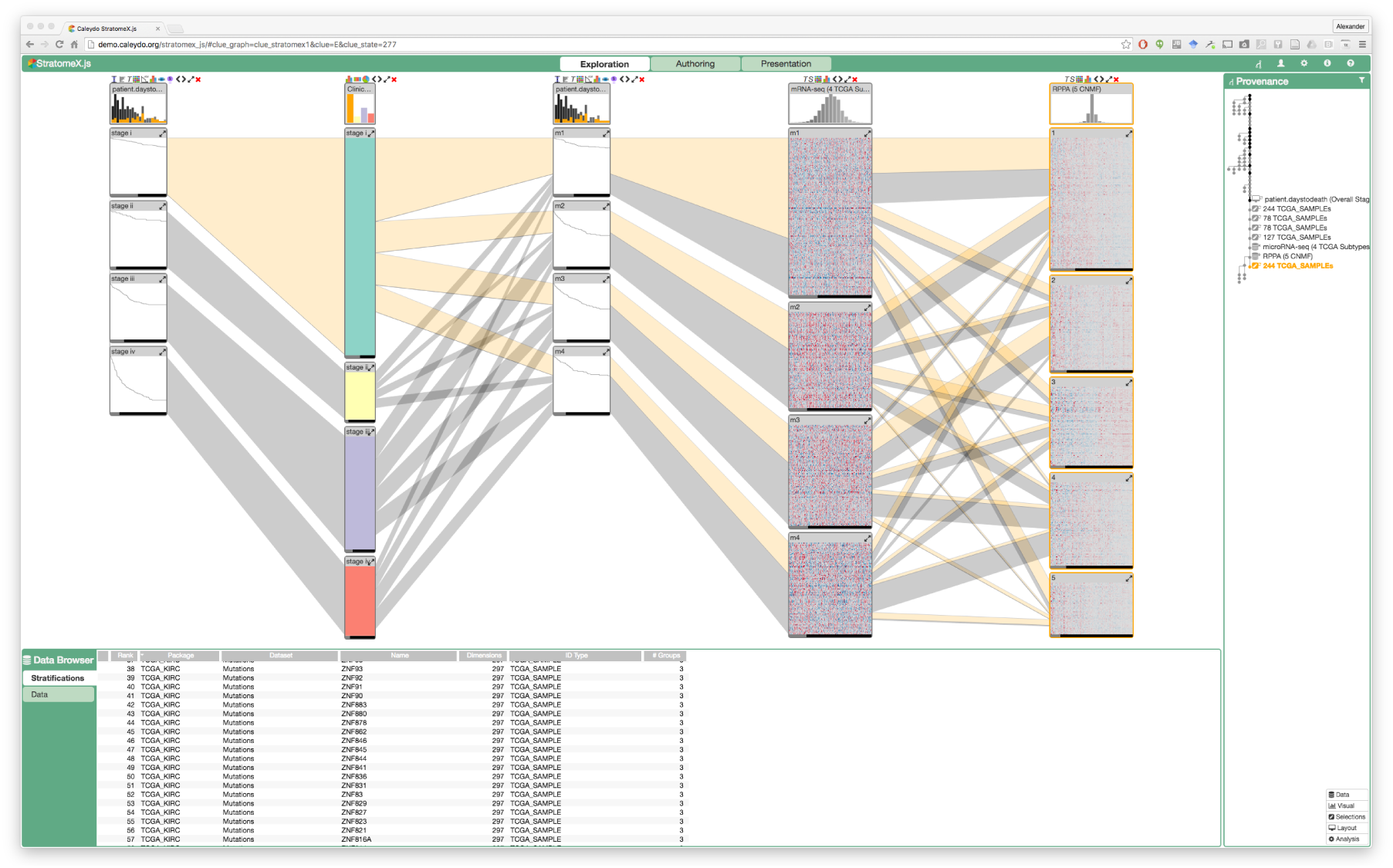
Screenshot of Caleydo StratomeX, which forms the basis of the technique introduced in this paper showing data from the TCGA Kidney Renal Clear Cell Carcinoma dataset [3]. Each column represents a dataset, which can either be categorical, like in the second column from the left, or based on the clustering of a high-dimensional dataset, like the two columns on the right. The blocks in the columns represent groups of records, where matrices are visualized as heat maps, categories with colors, and clinical data as Kaplan-Meier plots. The columns showing Kaplan-Meier plots are “dependent columns”, i.e., they use the same stratification as a neighboring column. The Kaplan-Meier plots show survival times from patients. The first column shows survival data stratified by tumor staging, where, as expected, higher tumor stages correlate with worse outcomes.

StratomeX visualizes stratifications of samples (patients) as rows (records) based on various attributes, such as clinical variables like gender or tumor staging, bins of numerical vectors, such as binned values of copy number alterations, or clusters of matrices/heat maps. Within these heat maps, the columns correspond to e.g., differentially expressed genes. StratomeX combines the visual metaphor used in parallel sets [19], with visualizations of the underlying data [20]. Each dataset is shown as a column. A header block at the top shows the distribution of the whole dataset, while groups of patients are shown as blocks in the columns. Relationships between blocks are visualized by ribbons whose thickness represents the number of patients shared across two bricks. This method can be used to visualize relationships between groupings and clusterings of different data, but can equally be used to compare multiple clusterings of the same dataset.

StratomeX also integrates the visualization of “dependent data” by using the stratification of a neighboring column for a different dataset. This is commonly used to visualize survival data in Kaplan-Meier plots for a particular stratification, for example, or to visualize expression of a patient cluster in a particular biological pathway.

## Related Work

There are several tools to analyze clustering results and assess the quality of clustering algorithms. A common approach to evaluate clustering results is to visualize the underlying data: heatmaps [1], for example, enable users to judge how consistent a pattern is within a cluster for high-dimensional data.

Seo at el. [21] introduced the hierarchical clustering explorer (HCE) to visualize hierarchical clustering results. It combines several visualization techniques such as scattergrams, histograms, heatmaps and dendrogram views. In addition to that, it supports dynamic partitioning of clusters by cutting the dendrogram interactively. HCE also enables the comparison of different clustering results while showing the relationship among two clusters with connecting links. Mayday [22, 23] is a similar tool that, in contrast to HCE, provides a wide variety of clustering options.

Lex et al. [24] introduced Matchmaker, a method that enables both, comparisons of clustering algorithms, and clustering and visualization of homogeneous subsets, with the intention of producing better clustering results. Matchmaker uses a hybrid heatmap and a parallel sets or parallel coordinates layout to show relationships between columns, similar to StratomeX. VisBricks [20] is an extension of this idea and provides multiform visualization for the data represented by clusters: users can choose which visualization technique to use for which cluster.

In contrast to these techniques, Domino [25] provides a completely flexible arrangement of data subsets that can be used to create a wide range of visual representations, including the Matchmaker representation. It is, however, less suitable for cluster evaluation and comparison.

A tool that addresses the interactive exploration of fuzzy clustering in combination with biclustering results is FURBY [26]. It uses a force-directed node-link layout, representing clusters as nodes and the relationship between them as links. The distance between nodes encodes the (approximate) similarity of two nodes. FURBY also allows users to refine or improve fuzzy clusterings by choosing a threshold that transforms fuzzy clusters into discrete ones.

A tool that is more closely related to our work is XCluSim [27]. It focuses on visual exploration and validation of different clustering algorithms and the concordance or disconcordance among them. It combines several small sub-views to form a multiview layout and provides different views for cluster evaluation. It contains dendrogram and force-directed graph views to show concordance among different clustering results and uses colors to represent clusters, without showing the underlying data. It lists used clustering algorithms, its parameters and applied similarity metrics. It offers a parallel sets view where each row represents one clustering result and thick dark ribbons depict which groups are clustered throughout all clustering results, referred as stable groups. In contrast to XCluSim, our methods integrate cluster metrics with the data more closely and can also bring in other, related data sources, to evaluate clusters. Also, XCluSim does not support cluster refinement.

Our methods are also related to silhouette plots, which visualize the tightness and separation of the elements in a cluster [28]. Silhouette plots, however, work best for geometric distances and clearly separated and spherical clusters, whereas our approach is more flexible in terms of supporting a variety of different measures of cluster fit. Also, silhouette plots are typically static, however, we could conceivably integrate the metrics used for silhouette plots in our approach. iGPSe [29], for example, is a system similar to StratomeX that integrates silhouette plots.

## Requirements

Based on our experience in designing multiple tools for visualizing clustered biomolecular data [24, 20, 11, 12, 30, 25], conversations with bioinformaticians, and a literature review, we elicited a list of requirements that a tool for the analysis of clustered, matrices from the biomolecular domain should address.

### R I: Provide representative algorithms with control over parametrization

A good cluster analysis tool should enable investigators to flexibly run various clustering algorithms on the data. Users should have control over all clustering parameters and should be able to choose from various similarity metrics.

### R II: Work with discrete, hierarchical and probabilistic cluster assignments

Visualization tools that deal with the analysis of cluster assignments should be able to work with all important types of clustering, namely discrete/partitional, hierarchical, and fuzzy clustering. The visualization of hierarchical and fuzzy clusterings is usually more challenging: to deal with hierarchical clusterings a tool needs to enable dendrogram cuts, and to address the properties of fuzzy clusterings, it must support the analysis of ambiguous and/or redundant assignments.

### R III: Enable comparison of cluster assignments

Given the ability to run multiple clustering algorithms, it is essential to enable the comparison of the clustering results. This will allow analysts to judge similarities and differences between algorithms, parametrizations, and similarity measures. It will also enable them to identify stable clusters, i.e., those that are robust to changes in parameters and algorithms.

### R IV: Visualize fit of records to their cluster

For the assessment of confidence in cluster assignments, a tool should show the quality of cluster assignments for its records and the overall quality for the cluster. This enables analysts to judge whether a record is a good fit to a cluster or whether it’s an outlier or a bad fit.

### R V: Visualize fit of records to other clusters

Clustering algorithms commonly don’t find the perfect fit for a record. Hence, it is useful to enable analysts to investigate if particular records are good fits for other clusters, or whether they are very specific to their assigned clusters. This allows users to consider whether records should be moved to other clusters, whether a group of records should be split off into a separate cluster, and more generally, to evaluate whether the number of clusters in a clustering result is correct.

### R VI: Enable refinement of clusters

To enable the improvement of clusters, users should be able to interactively modify clusters. This includes shifting of elements to better fitting clusters based on similarity, merging clusters considered to be similar, and excluding non-fitting groups from individual groups or the whole dataset.

### R VII: Visualize context for clusters

It is important to explore evidence for clusters in other data sources. In molecular biology applications in particular, datasets rarely stand alone but are connected to a wealth of other (meta)data. Judging clusters based on effects in other data sources can indicate practical relevance of a clustering, or can reveal dependencies between data sets and hence is important for validation and interpretation of the results.

## Design

We designed our methods to address the aforementioned requirements while taking into account usability and good visualization design practices. Our design was influenced by our decision to integrate the methods into Caleydo StratomeX as StratomeX is a well-established tool for subtype analysis. A prototype of our methods is available at goo.gl/I2qOxz. Please also refer to the supplementary video for an introduction and to observe the interaction.

We developed a model workflow for the analysis and refinement of clustered data, illustrated in Figure 2. This workflow is made up of four core components: (1) running a clustering algorithm, (2) visual exploration of the results, (3) manual refinement of the clustering results, and (4) interpretation of the results.

**Figure 2.**
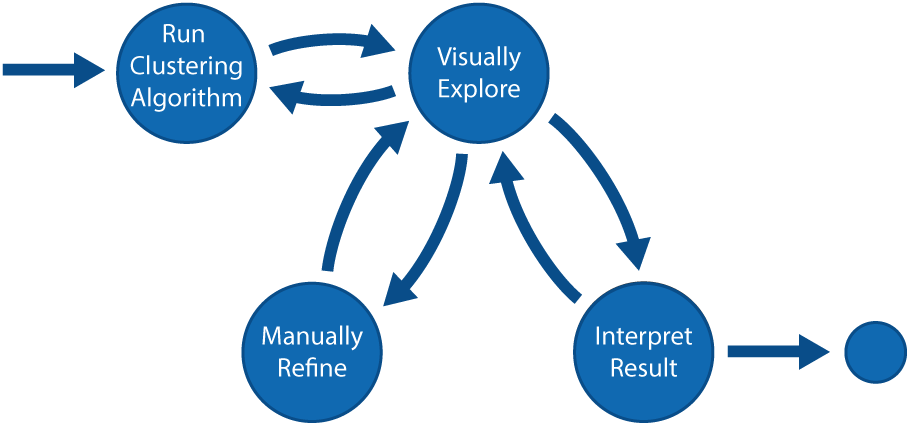
The workflow for evaluating and refining cluster assignments: (1) running clustering algorithms, (2) visual exploration of clustering results by investigating cluster quality and comparing cluster results (3) manual refinement and improvement of unreliable clusters and (4) final interpretation of the improved results considering contextual data.

### 1. Cluster Creation

Investigators start by choosing a dataset and either applying clustering algorithms with desired parametrization or selecting existing, precomputed clustering results. The clustered dataset is added to potentially already existing datasets and clusterings.

### 2. Visual Exploration

Once a dataset and clustering are chosen, analysts explore the consistency of clusters and/or compare the results to other clustering outcomes to discover patterns, outliers or ambiguities. If there are not confident about the quality of the result, or want to see an alternative clusterings, they can return to step 1 and create new clusters by adjusting the parameters or selecting a different algorithm.

### 3. Manual Refinement

If analysts detect records that are ambiguous, they can manually improve clusters to create better stratifications in a process that iterates between refinement and exploration. The refinement process includes splitting, merging and removing of clusters.

### 4. Result Interpretation

Once clusters are found to be of reasonable quality, the analysts can proceed to interpret the results. In the case of disease subtype analysis with StratomeX, they can assess the clinical relevance of subtypes, or explore relationships to other genomic datasets, confounding factors, etc. Of course, supplemental data can also inform the exploration and refinement steps.

We now introduce a set of techniques to address our proposed requirements within this workflow.

### Creating Clusters

Users are able to create clusters by selecting a dataset from a data browser window and choosing an algorithm and its configuration (see Figure 3). In our prototype, we provide a selection of commonly used algorithms in bioinformatics, including k-Means, (agglomerative) hierarchical clustering, Affinity Propagation, and Fuzzy c-Means. Each row in the popup represents one clustering technique with corresponding parameters, such as the number of clusters for k-Means, the similarity metric for hierarchical clustering, or the fuzziness factor for Fuzzy c-Means, addressing R I. Each execution of a clustering algorithm adds a new column of data to StratomeX, so that multiple alternative results can be easily compared.

**Figure 3.**
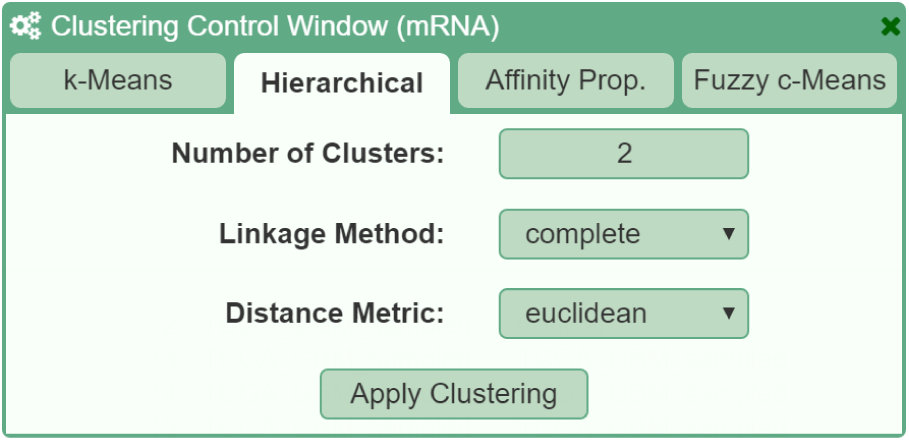
Example of the control window to apply clustering algorithms on mRNA genomic data. Different algorithms are accessible using tabs. Within the tabs, the algorithm can be configured using algorithm-specific parameters and general distance metrics.

### Cluster Evaluation

In our application, there are two components that enable analysts to evaluate cluster assignments: 1) the display of the underlying data in heatmaps or other visualizations and 2) the visualizations of cluster fit alongside the heatmap, as illustrated in Figure 4. The cluster fit data is either a measure of similarity of each record to the cluster centroid, or, if fuzzy clustering is used, the measure of probability that a record belongs to a cluster. Combining heatmaps and distance data allows users to relate patterns or conspicuous groups in the heatmap to their measure of fit.

**Figure 4.**
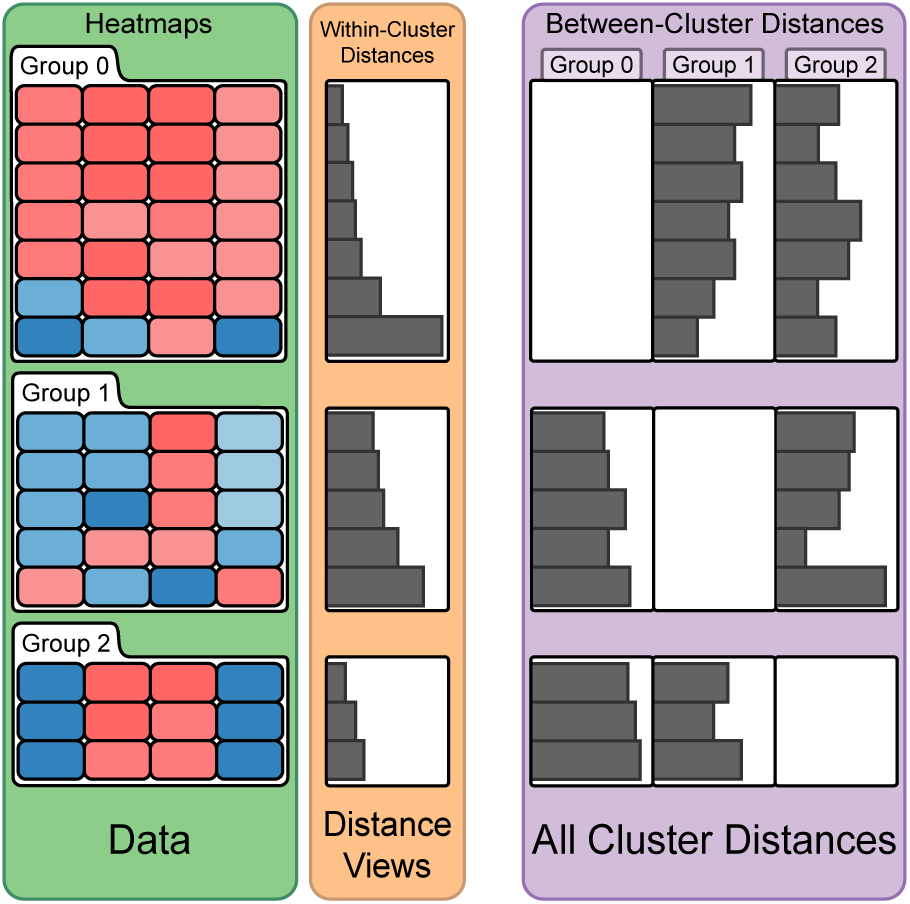
Illustration of heatmaps, within-cluster and between-cluster distance views. The within-cluster distance view shows the quality of fit of each record to its cluster (orange). The between-cluster distance view shows the quality of fit of each record to each other cluster (violet).

To evaluate the fit of each record to its cluster (R IV), we use a distance view, right next to the heatmap (orange in Figure 4). It displays a bar-chart showing the distances of each record to the cluster centroid. Each bar is aligned with the rows in the heatmap and thus represents the distance or correlation value of the corresponding record to the cluster mean. The length of a bar encodes the distance, meaning that short bars indicate well fitting records while long bars indicate records that are a poor fit. In the case of crosscorrelation, long bars represent records with high concordance whereas small bars indicate a disconcordance among them. While the absolute values of distances are typically not relevant for judging the fit of elements to the cluster, we show them on mouse-over in a tool-tip. The heatmaps and distance views are automatically sorted from best to worst fit which makes identifying the overall quality of a cluster easy. In addition to that, we globally scaled the length of each bar according to its distance measure, so that the largest bar represents the maximal computed distance measure across all distance views while still fitting into the distance view. Note that the distance measure used for the distance view does not have to be the one that was used for clustering. Figure 5 shows a montage of different distance measures for the same cluster in distance views. Notice that while some trends are consistent across many measures here, this is not for all measures and all patterns, illustrating the strong influence of the choice of a similarity measure.

**Figure 5.**
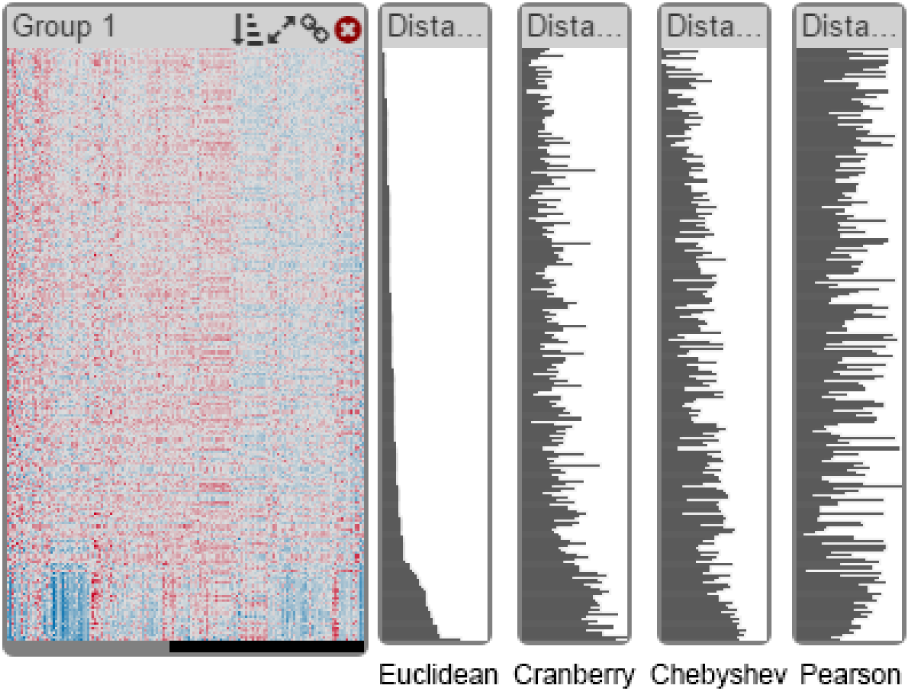
A montage of distance views showing different distance metrics for the same cluster.

Related to cluster fit is the question about the specificity of a record to a cluster (R V). It is conceivable that a record is a fit to multiple clusters, or that it would be a better fit to another cluster. To convey this, we compute the distances of each record to all other cluster centroids and visualize it in a **matrix of distances** to the right of the within-cluster distance view (violet in Figure 4). In doing so, we keep the row associations intact. We do not display the within-cluster distances in the matrix, which results in empty cells along the diagonal. This view helps analysts to investigate ambiguous records and supports them in judging whether the number of clusters is correct: if a lot of records have high distances to all clusters, maybe they should belong to a separate cluster. On demand, the heatmaps can also be sorted by any column in the between-cluster distance matrix. As an alternative to the bar charts, we also provide a grayscale heat map for between-cluster distances (see Figure 6), which scales better when the algorithm produced many clusters.

**Figure 6.**
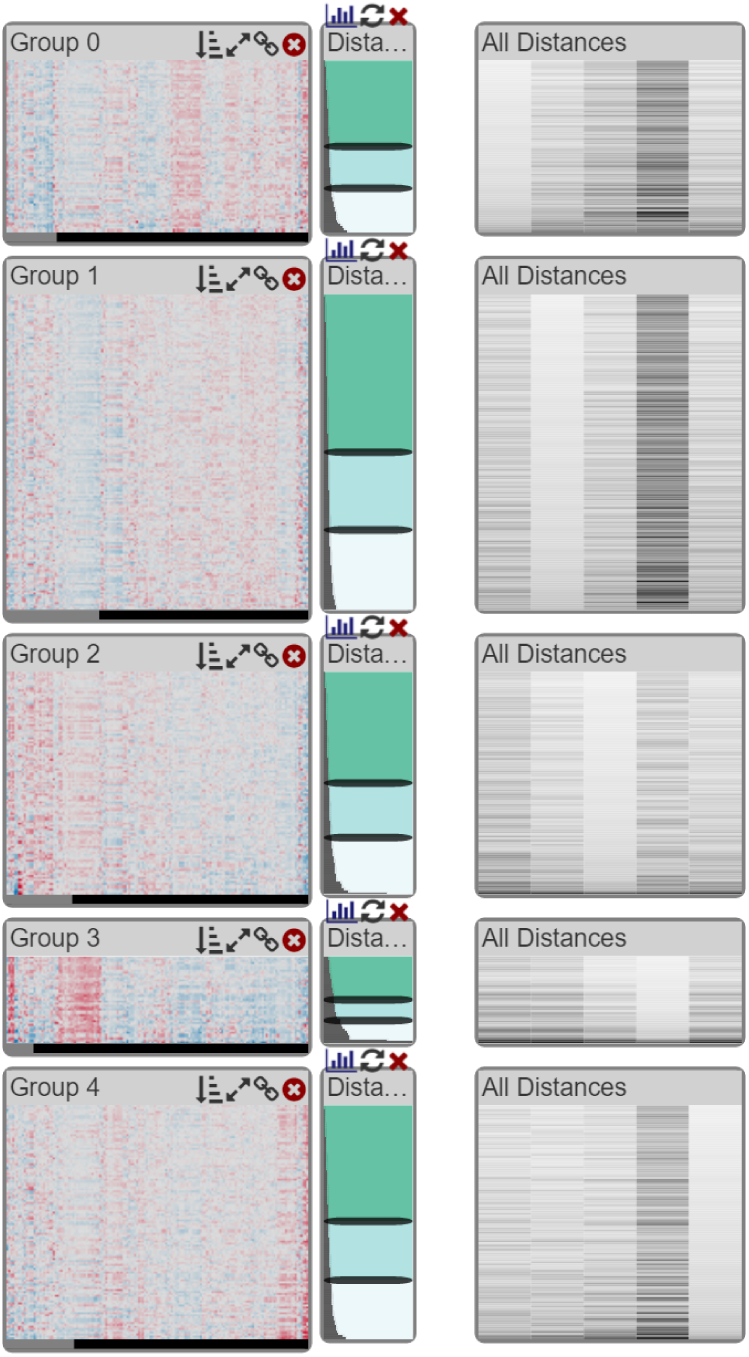
Example of five clusters, shown in heat maps, within-cluster distances shown to its right as bar charts, and between cluster distances on the far right as grayscale heat maps.

#### Visualizing Probabilities for Fuzzy Clustering

Since our tool also supports fuzzy clustering (R II) we provide a probability view, similar to the distance view, to show the degree of membership of each record to all clusters. In the **probability view**, the bars show the probability of a record belonging to a current cluster, which means that long bars always indicate a good fit. As each record has a certain probability to belong to each cluster, we use a threshold above which a record is displayed as a member of a cluster. Records can consequently occur in multiple clusters. Records that are assigned to multiple clusters are highlighted in purple, as shown in Figure 7, whereas unique records are shown in green. As for distance views, we also show probabilities of each record belonging to each cluster in a matrix, as shown in Figure 7 on the right.

**Figure 7.**
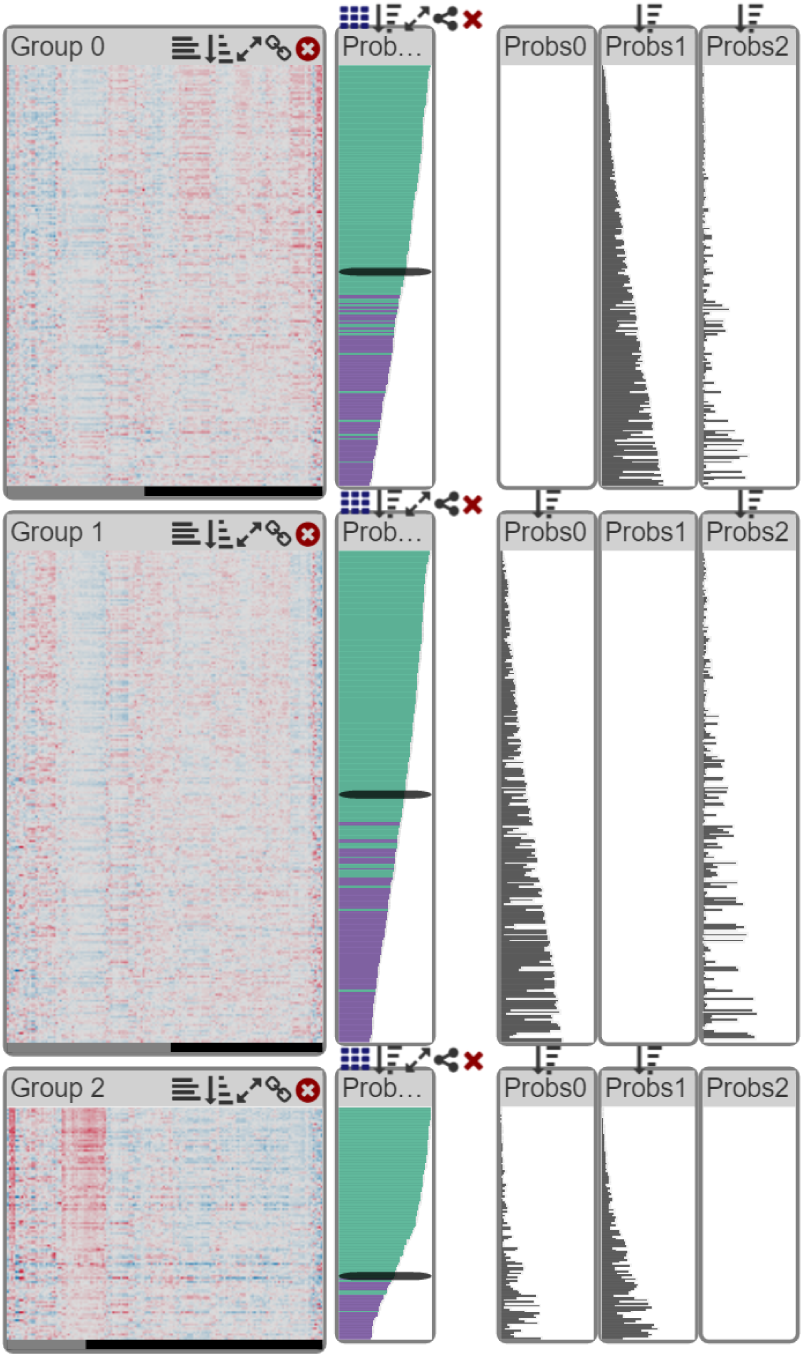
Example of three clusters produced by fuzzy clustering, shown in heatmaps. The probabilities of each patient belonging to their cluster are shown to their right. Green bars represent elements unique to the cluster while purple indicates patients belonging to more clusters. The between-cluster probabilities are displayed on the right.

### Cluster Refinement

Once scientists have explored the cluster assignments, the next step is to improve the cluster assignments if necessary (R VI).

#### Splitting Clusters

To support splitting of clusters, we extended StratomeX to enable analysts to define ambiguous regions in a cluster. The distance views also contain adjustable sliders that enable analysts to define up to three regions, to classify records into good, ambiguous, and bad fit (the green, light-green, and bright regions in Figure 8). By default, the sliders are set to the second and third quartile of the within-cluster distance distribution. Based on these definitions, analysts can split the cluster, which extracts the blocks into a separate column in StratomeX, as illustrated in Figure 8). This new column is treated like a dataset in its own right, such that the distance views show the distances to the new centroids, and clinical data can be interactively stratified based on these new blocks. However, these splits are not static: it is possible to dynamically adjust both sliders and hence the corresponding cluster subsets. In the context of fuzzy clustering, clusters can also be split based on probabilities. Clusters that were split and evaluated can then be re-integrated in the source column.

**Figure 8.**
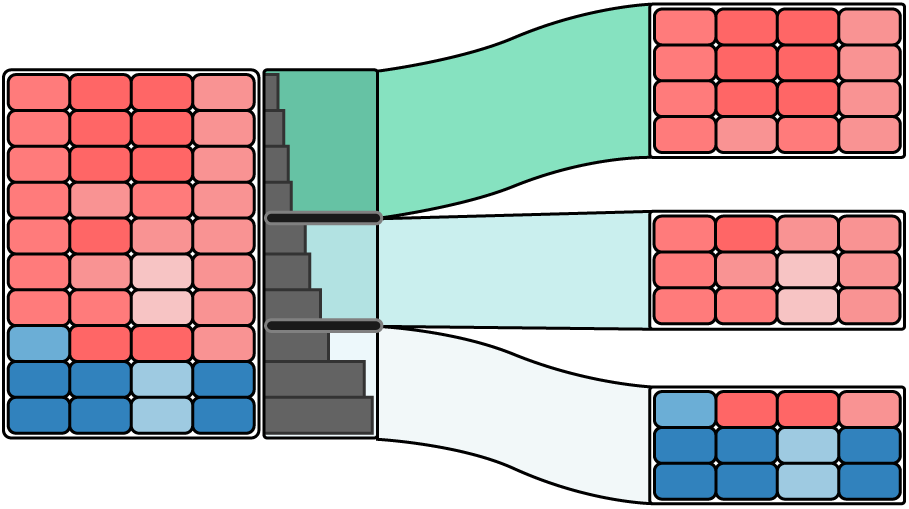
Example of a cluster division into three different subsets determined by two black sliders: (1) reliable, well fitting groups of elements (greenish color), (2) uncertain groups (blueish) and (3) unreliable, non-fitting elements (white color). The divisions are shown as separate column and related to the original cluster by colored ribbons.

#### Refining Split Clusters

Splitting just on distances does not guarantee that the resulting groups are as homogeneous as they could be: all they have in common is a certain distance range from the original centroid. To improve the homogeneity of split clusters, we can dynamically shift the elements between the clusters, so that the elements are in the cluster that is closest to them, based on the k-Means algorithm. In the future, we also plan on integrating re-clustering of individual blocks using arbitrary algorithms following this example.

#### Merging and Exclusion

Our application also has the option to merge clusters. Especially when several clusters are split first, it is likely that some of the new clusters exhibit a similar pattern, and that their distances also indicate that they could belong together, which can be addressed with a merge. We also support cluster exclusion since there might be groups or individual records that are outliers.

### Integration with StratomeX

The original StratomeX technique already enables cluster comparison R III through the columns and ribbons approach. It also is instrumental in bringing in contextual information for clusters R VII, as mentioned before. This can, for example, be used to asses the impact of refined clusterings on phenotypes. Figure 9 shows the impact of a cluster split on survival data, for example.

**Figure 9.**
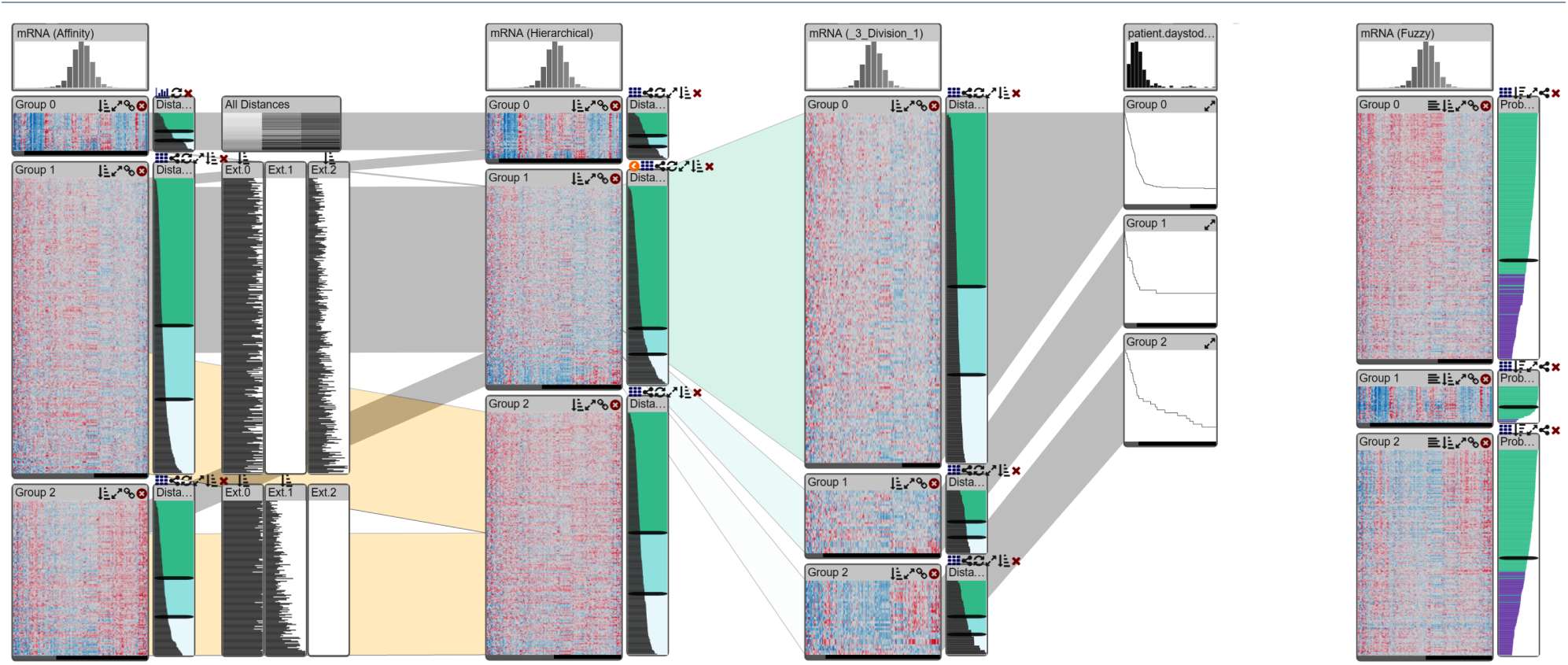
Overview of the improved StratomeX. The first column is stratified into three groups using affinity propagation. Distances between all clusters are shown. The second column shows the same data but is clustered differently using a hierarchical algorithm. Notice that Group 2 in the second column is a combination of Group 1 and Group 2 of the first column. The second block (Group 1) of the second column is split, and we see clearly that the patterns in the block at the bottom is quite different from the others. This block also exhibits a slightly differing phenotype: the Kaplan-Meier plot shows worse outcomes for this block. The rightmost column shows the same dataset clustered with a fuzzy algorithm. Notice that the second cluster contains mostly unique records (most bars are green).

## Usage Scenario

A common question in clustering is how to determine the appropriate number of clusters in the data. While there are algorithmic approaches, such as the cophenetic correlation coefficient [31], to estimate the number of clusters, visual inspection is often the initial step in confirming that a clustering algorithm has separated the elements appropriately. In this usage scenario we use our approach to inspect and refine a clustering result provided by an external clustering algorithm and to confirm our results with an integrated clustering algorithm.

We obtained mRNA gene expression data from the glioblastoma multiforme cohort of The Cancer Genome Atlas study [2] as well as clustering results generated using a consensus non-negative matrix factorization (CNMF) [32]. Verhaak et al. [2] reported four expression-derived gene expression subtypes in glioblastoma, which motivated us to review the automatically generated, uncurated, CNMF clustering results with 4 clusters. Visual inspection indicates that clusters named Group 0 and Group 1 contain patients (see Figure 10, leftmost column) that appear to have expression profiles that are very different from the other patients. Using the within-cluster distance visualization and sorting the patients in those clusters according to the within-cluster distance reveals that the expression patterns are indeed very different and that the within-cluster distances for those patients are also notably larger than for the other patients. Resorting the clusters by between-cluster distances to the other 3 clusters, respectively, shows that these patients are also different from the patients in the other clusters (see Figure 10).

**Figure 10.**
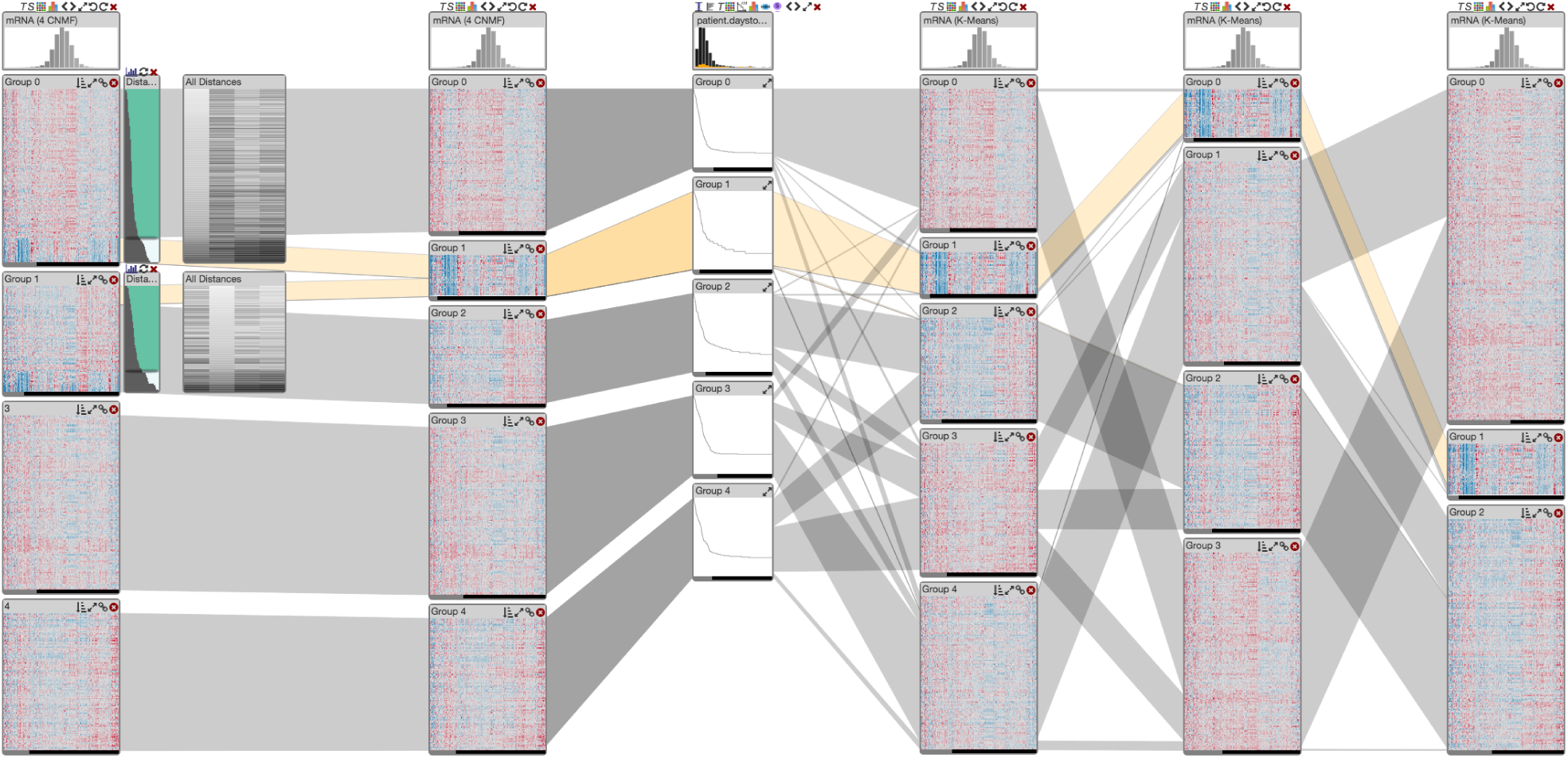
Results of manual cluster refinement and comparison with additional data types and clustering results. The original clustering is shown in the leftmost column. After sorting by within-cluster distances, the bottom sections of the corresponding blocks contain the expression profiles with poor cluster fit. The second column from the left shows the manually refined clustering with 5 clusters. The Kaplan-Meier plots show patient survival for each of the 5 clusters in the second column. The three rightmost columns are k-Means clustering results computed with the software. The derived cluster in the second column from the left is highlighted and the orange bands indicate overlap with patients in the selected cluster.

Using the sliders in the within-cluster distance visualization and the cluster splitting function we separated aforementioned patients from the clusters named Group 0 and Group 1. Because their profiles are very similar, we merged them into a single cluster using the cluster merging function. The expression profiles in the resulting new cluster look homogeneous and are visibly different from the expression profiles in the other 4 clusters. We examined patient survival times and age a disease diagnosis across the 5 clusters and did not observe any notable differences in the new cluster (see Figure 10). Since the web-based prototype of StratomeX is currently still lacking the guided exploration features of the original standalone application [12], we were unable to identify a meaningful correlation between the new cluster and mutation and copy number calls or to identify significantly overlapping clusters in other data types.

However, we also compared the five clusters derived from the original 4-cluster CNMF result with other clustering results computed on the same gene expression matrix and found, for example, that 3-, 4-, and 5-cluster k-Means clustering results using Euclidean distance and the k-means algorithm include almost exactly the same cluster that we identified in the CNMF clustering results using visual inspection and manual refinement.

## Implementation

Our tools are fully integrated with the web-version of Caleydo StratomeX. The clustering algorithms and distance computation are implemented as server plugins for Caleydo Web [33] in Python. We use the algorithms as implemented in the SciPy and the NumPy libraries. The front end is implemented as a client-side Caleydo web plugin and uses JavaScript and D3 [34]. The source code is released under the BSD license and is available at https://github.com/K3rn1n4tor/stratomex_js.

## Scalability and Limitations

Our methods are limited by the inherent limitations of StratomeX: when working with a large number of clusters, ribbons between the individual columns can create clutter. We observe that 10-15 clusters can be used without too much clutter. Also, the number of columns is limited to about 10 on typical displays. In terms of computational scalability, we found that even the computationally complex clustering algorithms such as affinity propagation execute almost interactively for a dataset with about 500x1500 entries, and complete within one to two minutes for a genomic dataset with about 500x12000 entries on our *t2.micro Amazon EC2 instance* with 1 CPU and 1 GB memory.

Our implementation currently cannot appropriately compare columns clustered with fuzzy algorithms, as the ribbons connecting the columns assume that every row exists only once. We plan on addressing this limitation in the future. Also, there are certain minor improvements of the user interface, for example, for the cluster dialog, that we intend to implement.

## Discussion and Conclusions

Clustering is an important yet inherently imperfect process. In this paper we have introduced methods to evaluate and refine clustering algorithms for the application to matrix data, as it is commonly used in molecular biology. In contrast to previous approaches, we combine visualization of the data directly with visualization refinement of clusters while associating the updated of cluster quality. We also allow interactive refinement of clusters while associating the updated clusters with contextual data, which allows analysts to judge clusters not only by the data used for clustering, but also based on effects observable in related datasets.

In the future, we plan on extending our work to datasets that are not in matrix form. This will require novel visual representations, as there is no equivalent to the well-defined borders of cluster blocks when clustering graphs or textual data.

## Acknowledgements

We thank Samuel Gratzl for his help with the implementation. This work was funded in part by the US National Institutes of Health (U01 CA198935, P41 GM103545-17, R00 HG007583) and supported by a fellowship of the FITweltweit program of the German Academic Exchange Service (DAAD).

## List of abbreviations used

*GBM*: glioblastoma multiforme
*TCGA*: *The Cancer Genome Atlas*

## Competing interests

The authors declare that they have no competing interests.

